# Whole-Genome Characterization of *Proteus mirabilis* Isolated from Fermented Soybean, *hawaijar*: Insights into Foodborne Virulence and Antimicrobial Resistance Determinants

**DOI:** 10.64898/2026.01.09.698623

**Authors:** Moirangthem Goutam Singh, Romi Wahengbam

## Abstract

*Proteus mirabilis* is a well-known opportunistic pathogen primarily associated with urinary tract and wound infections in humans; however, its presence in fermented foods has rarely been documented. This study reports the first whole-genome characterization of P. mirabilis strain FFPR3MR5, isolated from traditionally fermented soybean (hawaijar) produced in Moirang, Manipur, India, to evaluate its genomic features, virulence potential, and evolutionary relationship with clinical isolates. The draft genome comprised 3.74 Mb with a GC content of 38.56%, assembled into 109 contigs and encoding 3,317 coding sequences, with 100% completeness and 0.17% contamination. Functional annotation identified 3,282 KEGG orthologs mapped to 40 metabolic pathways, with metabolism-related genes predominating (75.05%). Genome screening revealed 35 virulence-associated genes involved in adherence, motility, biofilm formation, and immune evasion, along with four antimicrobial resistance genes conferring resistance to β-lactams, fluoroquinolones, tetracyclines, and macrolides. Phylogenetic analysis showed that FFPR3MR5 clustered closely with the clinical reference strain *P. mirabilis* NC_010554, supported by a high OrthoANIu value of 99.39%. Genome-wide variant analysis identified approximately 27,000 polymorphisms, including deleterious nonsynonymous substitutions in genes associated with stress response (*cpxA, kdpD*), biofilm formation (*bcsA*), DNA repair (*recB, recC, ssb*), and metabolism (*metH*), suggesting niche-specific adaptation to the fermentation environment. The coexistence of virulence and antimicrobial resistance determinants in this food-derived strain underscores its potential as a reservoir of clinically relevant traits, highlighting the need for genomic surveillance of traditional fermented foods within a One Health framework.

**Importance:** This study offers a comprehensive whole-genome analysis of a *P. mirabilis* strain isolated from fermented soybean food, broadening current knowledge of how opportunistic pathogens survive outside clinical settings. Integrated genomic, phylogenetic, and variant analyses revealed the coexistence of virulence and antimicrobial resistance determinants, highlighting fermented foods as potential reservoirs of clinically relevant bacteria. Deleterious mutations in genes linked to biofilm formation, stress response, metabolism, and DNA repair suggest adaptation to fermentation environments, while close relatedness to a clinical reference strain underscores public health relevance. These findings emphasize the need for genomic surveillance of traditional fermented foods within a One Health framework.

## Introduction

Fermented soybeans are widely consumed in various Asian countries. For example, they are consumed in various forms, including *natto* (Japan), *tempeh* (Indonesia), *douchi* (China), *thua nao* (Thailand), and *choongkook jang* (Korea) (1). In India, the consumption of fermented soybeans is primarily found in the northeastern states. For instance, people in northeast India traditionally prepared and consumed a non-salted, sticky, fermented soybean product. This includes *kinema* of Sikkim, *hawaijar* of Manipur, *tungrymbai* of Meghalaya, *bekang* of Mizoram, *aakhone* of Nagaland, and *peruuyaan* of Arunachal Pradesh (2). Fermented soybeans offer several benefits, including immunomodulation, enhanced nutritional quality, antioxidant activity, and the production of bioactive peptides (3). Despite their health benefits, frequent outbreaks have been reported due to the consumption of fermented soybeans (4). This outbreak may be linked to the nature of production, the fermentation process, and market distribution. A study by Keisam *et al*. linked the risk of food poisoning and infection from consumption of fermented soybean due to two enteric bacterial pathogens *Bacillus cereus* and *Proteus mirabilis* (4).

*P. mirabilis* is a Gram-negative opportunistic pathogen that belongs to the family Enterobacteriaceae (5). It is widely present in the natural environment and the intestine of humans and animals. While *P. mirabilis* is a well-recognized cause of urinary tract and wound infections, its detection in food products suggests environmental adaptability and raises concerns about transmission through the food chain (6).

In recent years, whole-genome sequencing (WGS) has emerged as a transformative tool in food safety and microbial epidemiology. WGS provides the highest resolution for pathogen subtyping, enabling detailed characterization of virulence factors, antimicrobial resistance genes, and mobile genetic elements that classical methods cannot achieve. Guidance from the World Health Organization and numerous public health agencies highlights that WGS enhances routine surveillance, outbreak detection, source attribution, and response to foodborne diseases across the farm-to-fork continuum (7). WGS has now been integrated into national and international public health frameworks to monitor the emergence, spread, and evolution of foodborne pathogens with unprecedented precision (8).

Furthermore, genomic surveillance using WGS can detect the early emergence and dissemination of antimicrobial resistance (AMR), informing both clinical and food safety interventions. The WHO’s Global AMR Surveillance System (GLASS) now promotes WGS to track AMR determinants at local, national, and global scales, enabling timely policy development and targeted mitigation strategies (9). In parallel, extensive efforts such as the FDA’s GenomeTrakr program illustrate how WGS supports traceback investigations, facilitates outbreak source identification, and enhances the detection of emerging resistance threats in the food supply (10). These WGS-based approaches are indispensable for One Health strategies, enabling the integration of genomic data from human, animal, and environmental sources to capture the full ecological context of pathogen transmission and AMR spread (11).

In this study, we report the first whole-genome characterization of *P. mirabilis* strain FFPR3MR5 isolated from a fermented soybean (*hawaijar*) sample collected from Manipur, India. By integrating WGS, comparative genomics, and functional annotation, we elucidate its genomic features, virulence potential, antimicrobial resistance profile, and evolutionary relationship to clinical isolates. The findings provide insights into the genomic basis of pathogen adaptation to fermented food matrices and underscore the relevance of WGS in food safety and epidemiological surveillance.

## Results

### Identification of isolate

BLASTN analysis of the 16S rRNA gene sequence revealed *P. mirabilis* as the closest match. The query sequence showed 99.10–99.24% nucleotide identity with multiple *P. mirabilis* type strains, including *P. mirabilis* ATCC 29906 (GenBank accession NR_114419.1), with 100% query coverage and an E-value of 0.0, confirming species-level identification.

### Genome Overview

A total of 1,161,518 PE reads were trimmed, producing 989,927 high-quality (Q ≥ 30) PE reads. The genome size was 3,742,076 bp with 52-fold coverage, 38.56% GC content, and an N_50_ value of 155,540 bp with the largest contig length of 247,027 bp. It consisted of 109 contigs with 3,317 coding sequences, 4 rRNAs, 1 tmRNA, and 59 tRNAs. CheckM2 identified 100% completeness and 0.17% contamination (Fig 1).

**Fig 1.**
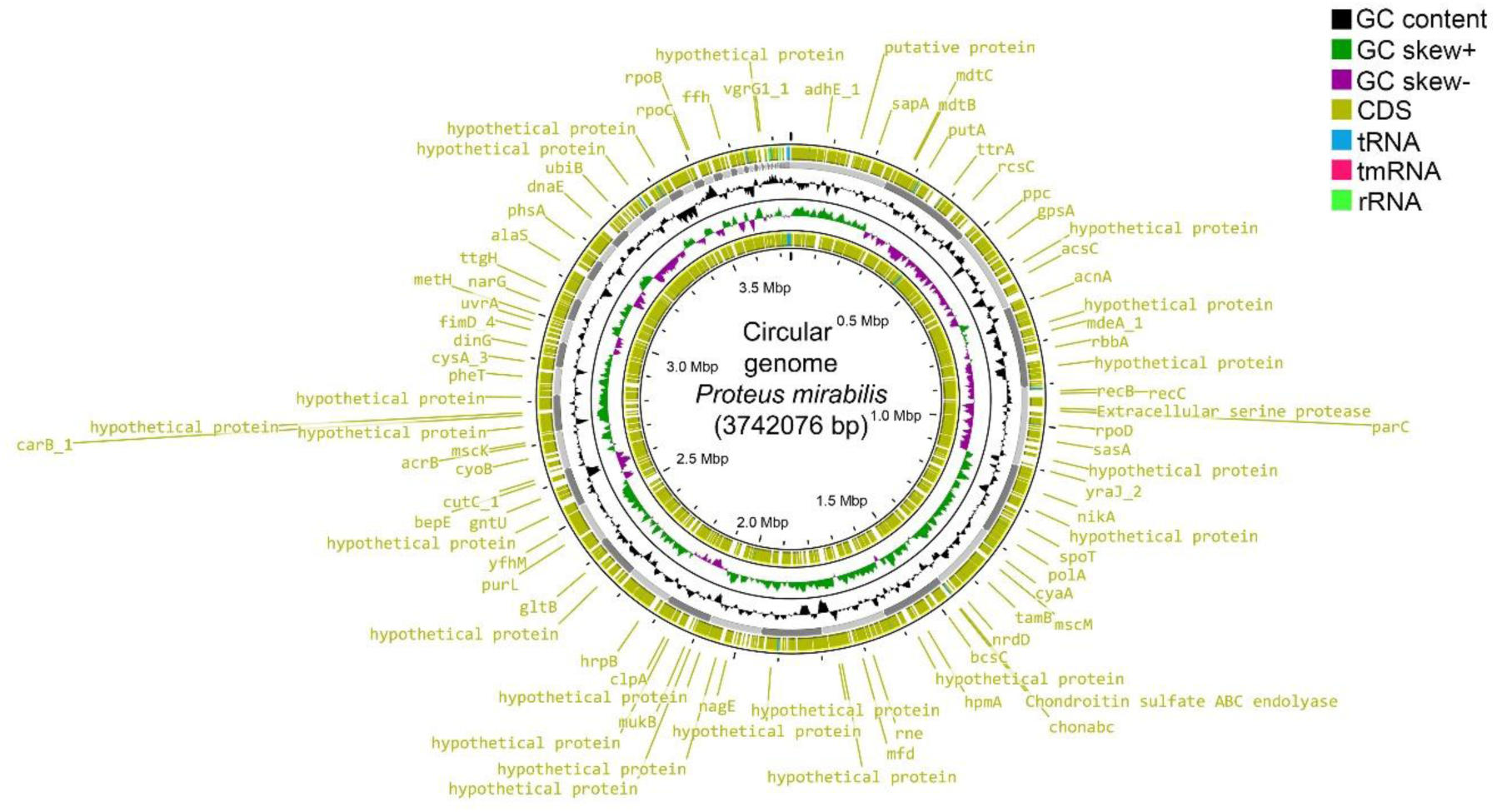
The draft circular genome of *Proteus mirabilis* (3,742,076 bp) is shown with annotated genomic features. From the outer to inner rings: predicted coding sequences (CDS) on the forward and reverse strands (yellow), tRNA genes (blue), tmRNA (pink), and rRNA operons (light green). The inner rings represent GC content (black) and GC skew, with positive skew shown in green and negative skew in purple. Selected genes of interest, including housekeeping genes, transporters, metabolic enzymes, and virulence-associated factors, are labeled around the chromosome. Genome coordinates are indicated in megabase pairs (Mbp). Only few genomes and their location are shown.

### Gene function analysis

Gene function annotation using databases such as Comprehensive Antibiotic Resistance Database (CARD), Virulence Factor Database (VFDB), Pathogen-Host Interaction (PHI), and Eggnog Mapper identified 4, 801, 356, 3282 (Kyoto Encyclopedia of Genes and Genomes (KEGG)) and 3047 (Cluster of Orthologs (COG)) genes, respectively. KEGG analysis revealed that 3,282 orthologous genes mapped to 40 metabolic pathways. The predominant categories were metabolism (75.05%), environmental Information processing (7.78%), genetic information processing (6.40%), cellular processes (5.09%), human diseases (4.27%), and organismal systems (1.43%) (Fig 2). The human disease category revealed a high abundance of genes associated with drug resistance (47) and infectious diseases (28) (Fig 2E). COG analysis annotated 3047 genes, categorized into 21 functional groups. Major functional groups and functional categories include transcription, energy production and conversion, translation, ribosomal structure and biogenesis, inorganic ion transport and mechanism, cell wall/membrane/envelope biogenesis, and coenzyme transport and mechanism (Fig 3A).

**Fig 2.**
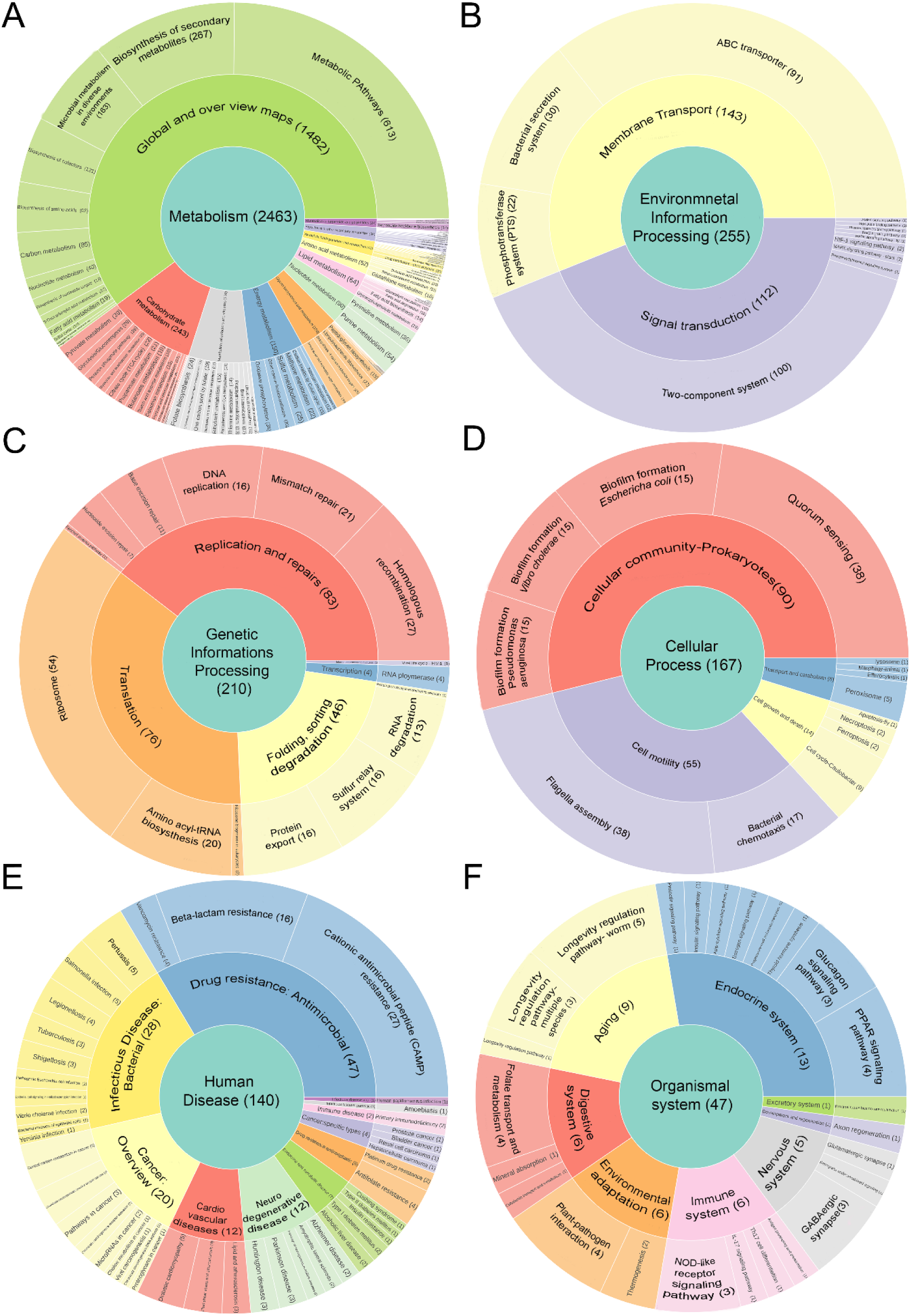
Functional annotation of the *Proteus mirabilis* genome based on KEGG pathway classification. Sunburst plots depict the hierarchical distribution of predicted genes across major KEGG functional categories and subcategories. Panels show: (A) Metabolism, (B) Environmental Information Processing, (C) Genetic Information Processing, (D) Cellular Processes, (E) Human Diseases, and (F) Organismal Systems. The inner circle represents the main KEGG category, while outer rings indicate progressively finer functional subdivisions.

**Fig 3.**
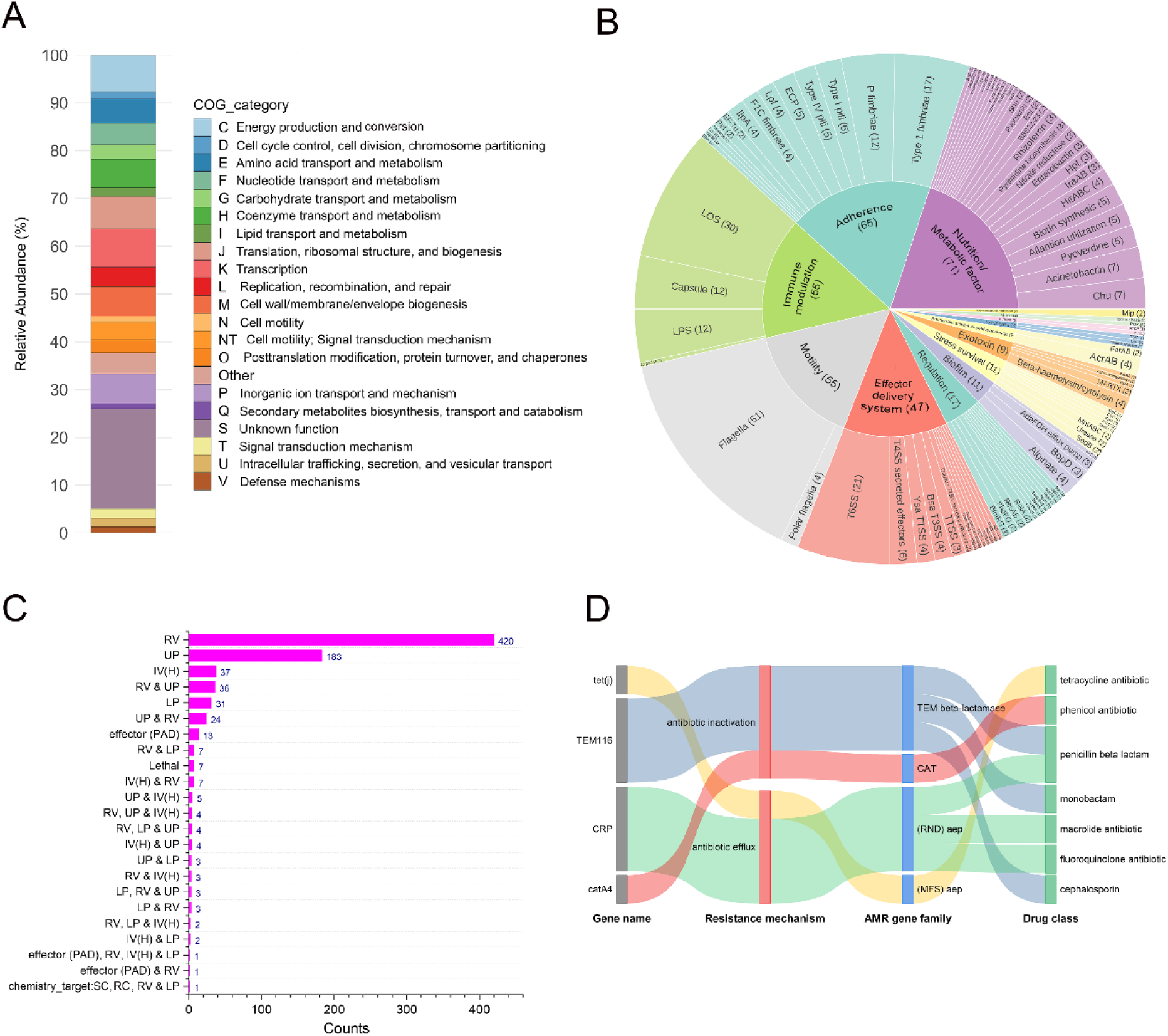
Functional classification, virulence factors, and antimicrobial resistance profile of Proteus mirabilis. (A) Distribution of predicted proteins across Clusters of Orthologous Groups (COG) functional categories, shown as relative abundance (%) (B) Sunburst plot of VFDB analysis classified by functional modules, including adherence, immune modulation, motility, nutrient acquisition, secretion systems, toxins, stress survival, and regulation; numbers indicate gene counts within each class. (C) Bar plot summarizing the number of virulence-associated genes grouped by functional roles and their combinations. Abbreviation: RV, reduced virulence; UP, unaffected pathogenicity; IV, increase virulence; H, hypervirulence; PAD, plant a virulence determinant; LP, loss of pathogenicity; SC, sensitivity to chemicals; RC, resistance to chemical. (D) Sankey diagram linking antimicrobial resistance (AMR) genes to resistance mechanisms, AMR gene families, and corresponding drug classes, illustrating the multidrug resistance potential of the genome. Abbreviation: tet, tetracycline; TEM, Temneira Enzyme Mutant; CRP, cyclic AMP receptor protein; RND, resistance-nodulation cell division; MFS, major facilitator superfamily; CAT, chloramphenicol acetyltransferase.

**Fig 4.**
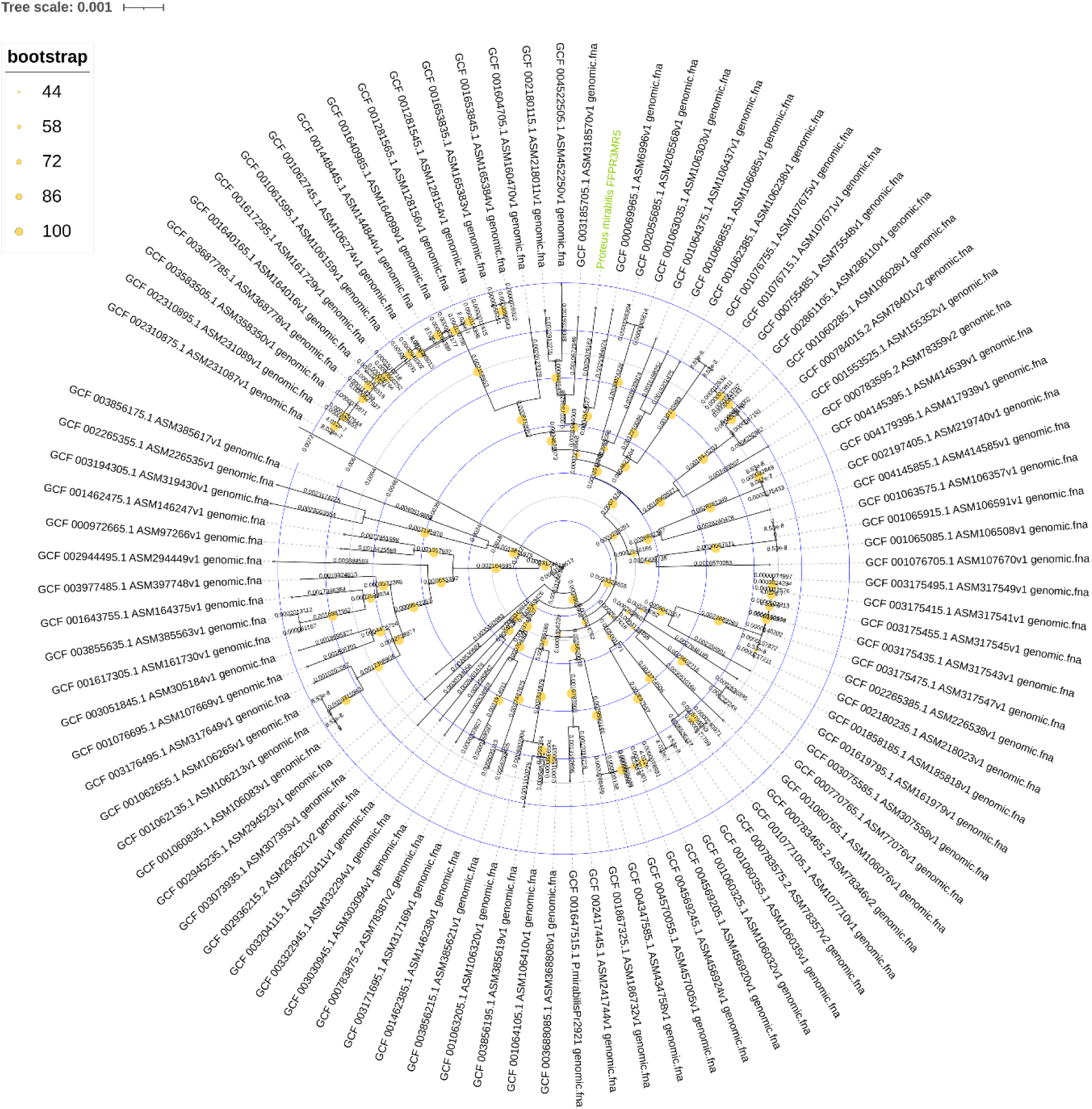
Circular phylogenetic tree of *Proteus mirabilis* strains based on whole-genome sequence analysis. The radial phylogeny depicts the evolutionary relationships among publicly available *P. mirabilis* genomes inferred from genome-wide comparisons. Branch lengths are proportional to genetic distance (scale bar = 0.001 substitutions per site). Bootstrap support was estimated using UltraFast bootstrap with 5,000 replicates and is shown as colored circles at internal nodes, with increasing intensity representing higher confidence levels (44–100%). The study strain is highlighted in green, illustrating its phylogenetic placement relative to reference and closely related *P. mirabilis* isolates.

Virulence factor analysis using the VFDB identified 35 putative virulence genes, with 15 genes having >80% identity (Table S1). These virulence genes were classified into 14 categories: nutritional/metabolic factor (19.94%), adherence (18.26%), motility (15.45%), immune modulation (15.45%), effector system (12.36%), regulation (3.93%), biofilm (3.09%), stress survival (3.09%), exotoxin (2.53%), exoenzyme (0.84%), invasion (0.56%), post transcriptional modification (0.56%), and other (0.56%) (Fig 3B). Notably, genes associated with flagella, lipooligosaccharide, type I fimbriae, capsule, lipopolysaccharide, and P fimbriae were the most abundant, which are critical for bacterial motility, adhesion to host tissues, immune evasion, biofilm formation, and overall virulence.

In the PHI database, 801 pathogen-host interaction-related genes were annotated (Table S2). These annotated genes were categorised into 23 groups, with reduced virulence (420 genes) being the most abundant, followed by unaffected pathogenicity (183 genes), increased virulence (Hypervirulence) (37 genes), loss of pathogenicity (31 genes), and lethal (7 genes) (Fig 3C).

In the CARD database, the strain FFPR3MR5 exhibited two antibiotic resistance mechanism: antibiotic inactivation and antibiotic efflux, covering 4 antibiotic resistance genes. All antibiotic resistance genes have an identity score greater than 80% (Table S3). The strain was resistant to tetracycline, phenicol, penicillin (beta-lactam), monobactam, macrolide, fluoroquinolone, and cephalosporin antibiotics (Fig. 3D).

### Phylogenetic Relationships

The final core genome alignment generated by Parsnp and exported with HarvestTools yielded a high-quality alignment suitable for phylogenetic reconstruction. Out of 102 genomes, 95 pass the core-genome alignment. Using the GTR+F+I+R7 model (BIC-selected) in IQ-TREE2, we constructed a robust maximum-likelihood phylogeny with 5,000 ultrafast bootstrap replicates. In iTOL, the bootstrap value is expressed as a range of 0-100, where 0 indicates no bootstrap, and 100 indicates 5000 bootstraps, based on the number of bootstraps applied in IQTREE. Bootstrap values ranged from moderate to very high (42–100), indicating strong support for most internal branches. The tree revealed distinct clustering patterns among the 95 *P. mirabilis* genomes, reflecting their evolutionary relationships and genomic similarity. Visualization in iTOL allowed detailed inspection of topologies, mapping of bootstrap support, and precise separation of phylogenetic clusters (Fig4). Phylogenetic analysis revealed that *P. mirabilis* FFPR3MR5 clustered tightly with the reference strain GCF_00069965.1 (NC_010554), separated by only a very short branch length, reflecting minimal nucleotide divergence. This relationship was further supported by the calculated OrthoANIu value of 99.39%, confirming that the two genomes share nearly identical nucleotide content across their aligned regions. ANI values above 95% are characteristic of the same species, while values exceeding 98–99% typically represent very closely related strains. Thus, the 99.39% ANI strongly indicates that FFPR3MR5 and GCF_00069965.1 belong to the same lineage and likely share similar biological traits.

### Genome-wide variant profiling

Comprehensive analysis of the VCF file revealed a total of ∼27,000 high-confidence variants distributed across the genome (Fig 5A). The majority of variants were located within protein-coding regions (13,732 variants) and annotated gene regions (13,506 variants), together accounting for nearly all detected variants. In contrast, very few variants were associated with non-coding RNA features, with only 21 variants in rRNA genes and 2 variants in tRNA genes, indicating strong enrichment of polymorphisms within coding sequences. Variant density plotted across genomic coordinates (per 1 kb windows) showed a heterogeneous distribution along the chromosome. While most regions exhibited moderate variant counts, several localized peaks were observed, indicating mutation hotspots or regions of elevated polymorphism (Fig 5B). No large variant-free regions were detected, suggesting genome-wide coverage of variants. Gene-level aggregation identified multiple genes harboring a high number of variants. Notably, genes involved in the redox reaction (*rsxC*), mobile genetic element (*tnpA*), metabolism (*metH* & *gltB*), chromosome organisation (*mukB*), and biofilm formation (*bcsA*) showed the highest variant counts, followed by genes involved in membrane integrity and transport (*yhdP* & *bamA*), DNA repair and homologous recombination (*recB*), hemolysin-associated cytotoxic activity (*hpmA*), and central carbon-nitrogen metabolism (*carB*). This pattern suggests that genetic variation is not randomly distributed, but instead enriched within specific functional categories (Fig 5C). Functional annotation revealed that synonymous variants were the most abundant class (21,600 variants), followed by nonsynonymous substitutions (5,570 variants) (Fig 5D). Variants predicted to have high-impact effects were relatively rare, including 73 frameshift and 18 nonsense mutations. A small number of variants were classified as NA (71), likely corresponding to features without definitive effect annotation. Overall, the dominance of synonymous and nonsynonymous variants indicates that most polymorphisms represent single-nucleotide substitutions with potentially moderate functional consequences.

**Fig 5.**
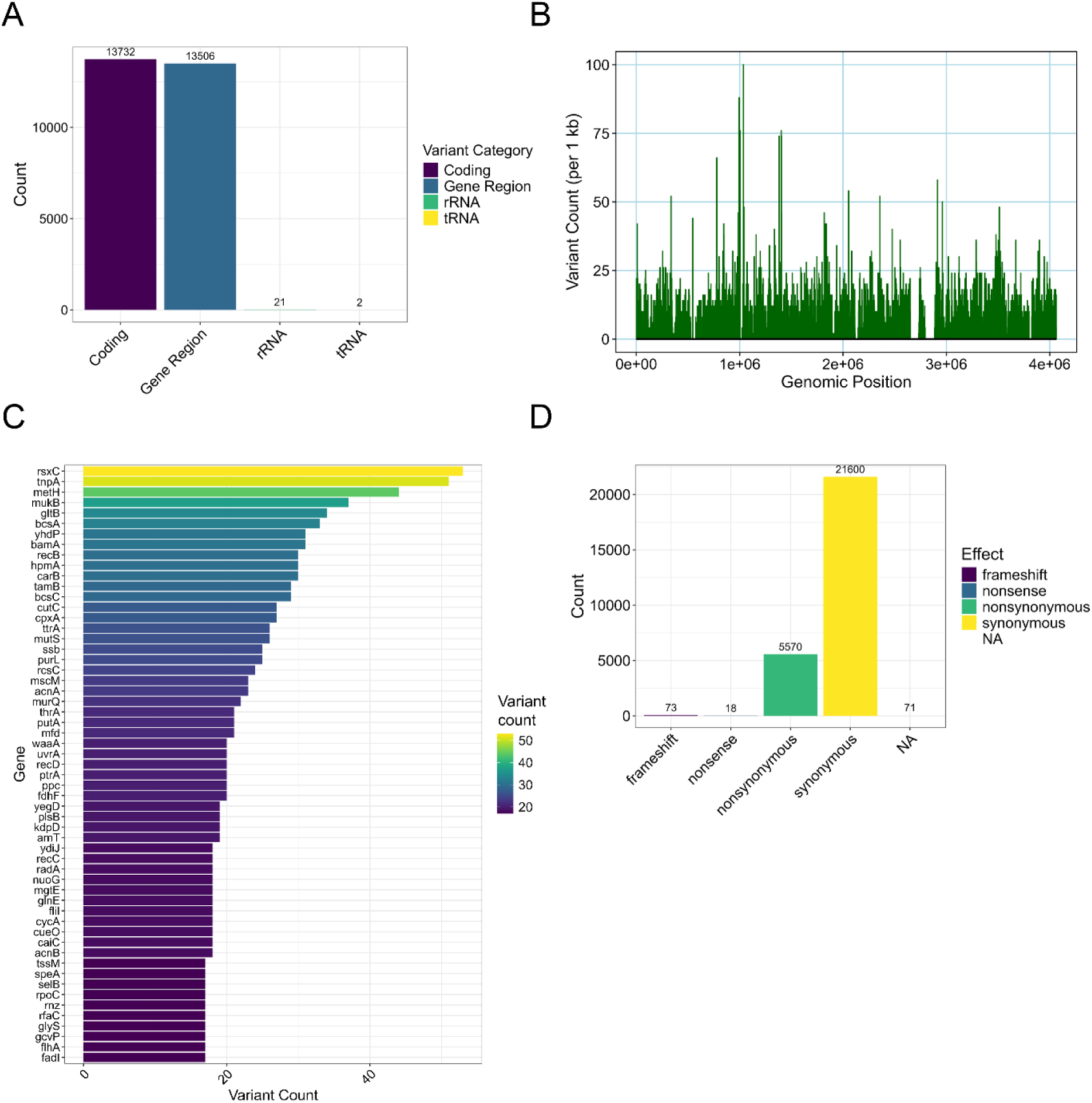
Genome-wide single-nucleotide variant (SNV) landscape of Proteus mirabilis. (A) Distribution of identified variants across genomic features, including coding regions, intergenic (gene) regions, rRNA, and tRNA genes (B) Genome-wide density of variants plotted as variant count per 1 kb window across chromosomal coordinates, highlighting regions with elevated polymorphism (C) Top genes harboring the highest number of variants, with bar color indicating variant frequency per gene (D) Functional classification of variants based on predicted effects, including synonymous, nonsynonymous, frameshift, nonsense, and unannotated (NA) variants, showing the predominance of synonymous substitutions.

Among the top 57 variant genes (Fig 5C), 45 genes had non-synonymous substitutions, and 12 genes had synonymous substitutions. Notably, deleterious variants (PROVEAN ≤ -2.5) were detected in genes such as *bcsA* (V701A), *cpxA* (P338S), *gcvP* (A159V), *kdpD* (A33T), *hpmA* (G1133D), *metH* (N794H), *recB* (A896V; R553S), *recC* (R112L), and *ssb* (P141S) with each of these genes harboring at least one predicted deleterious amino-acid substitution (Table 1). These genes are functionally diverse, contributing to processes including biofilm formation (bcsA), stress-response regulation (*cpxA*, *kdpD*), metabolism (*gcvP*, *metH*), virulence (*hpmA*), and DNA repair (*recB*, *recC*, *ssb*). The presence of deleterious variants in these loci suggests possible impairments in critical cellular pathways, including envelope stress sensing, metabolic regulation, and genomic maintenance. Collectively, these mutations may influence bacterial adaptability, virulence, and stress tolerance, indicating that functional alterations in these key genes could shape the strain’s physiological and pathogenic characteristics.

**Table 1.**
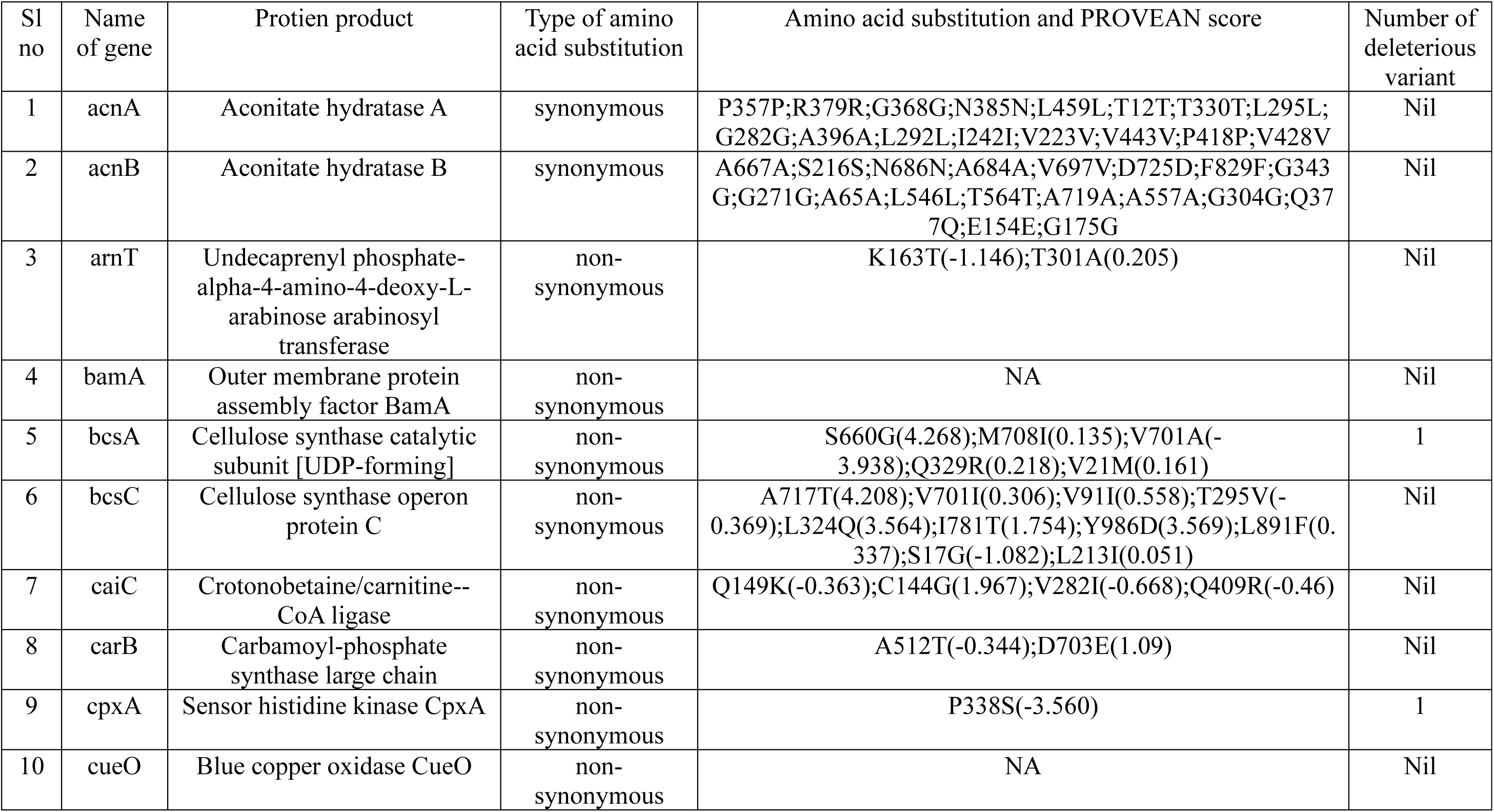
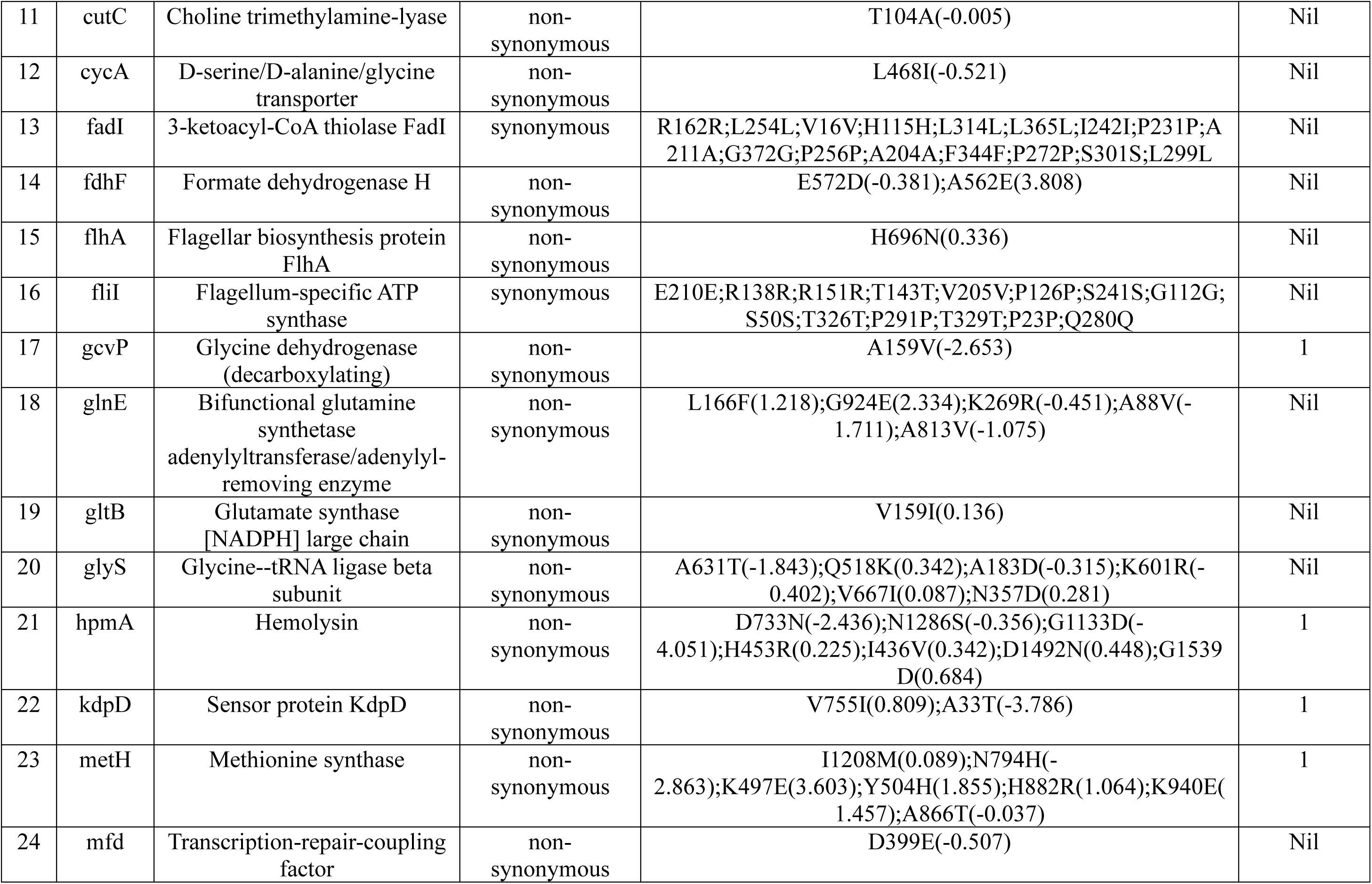
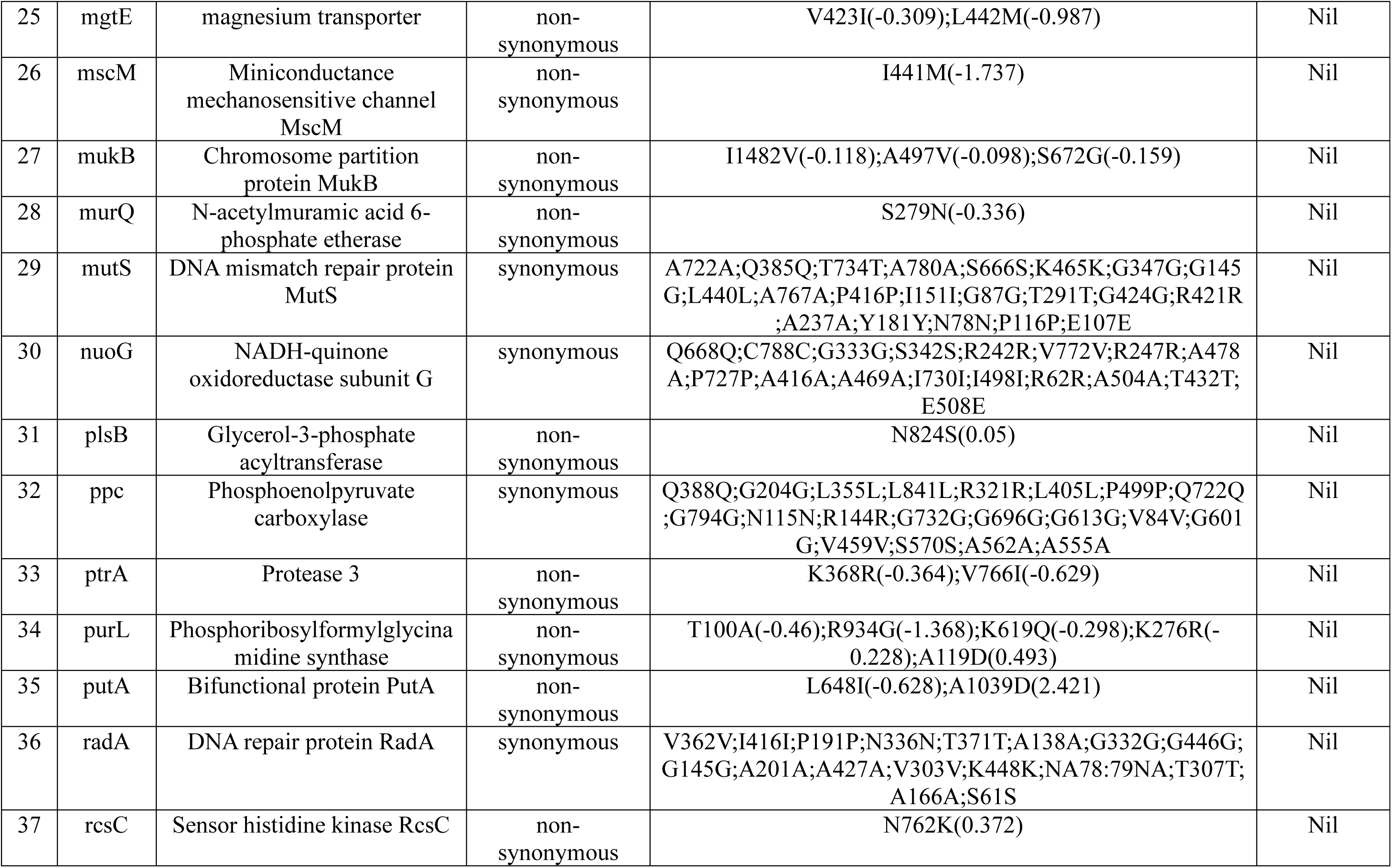
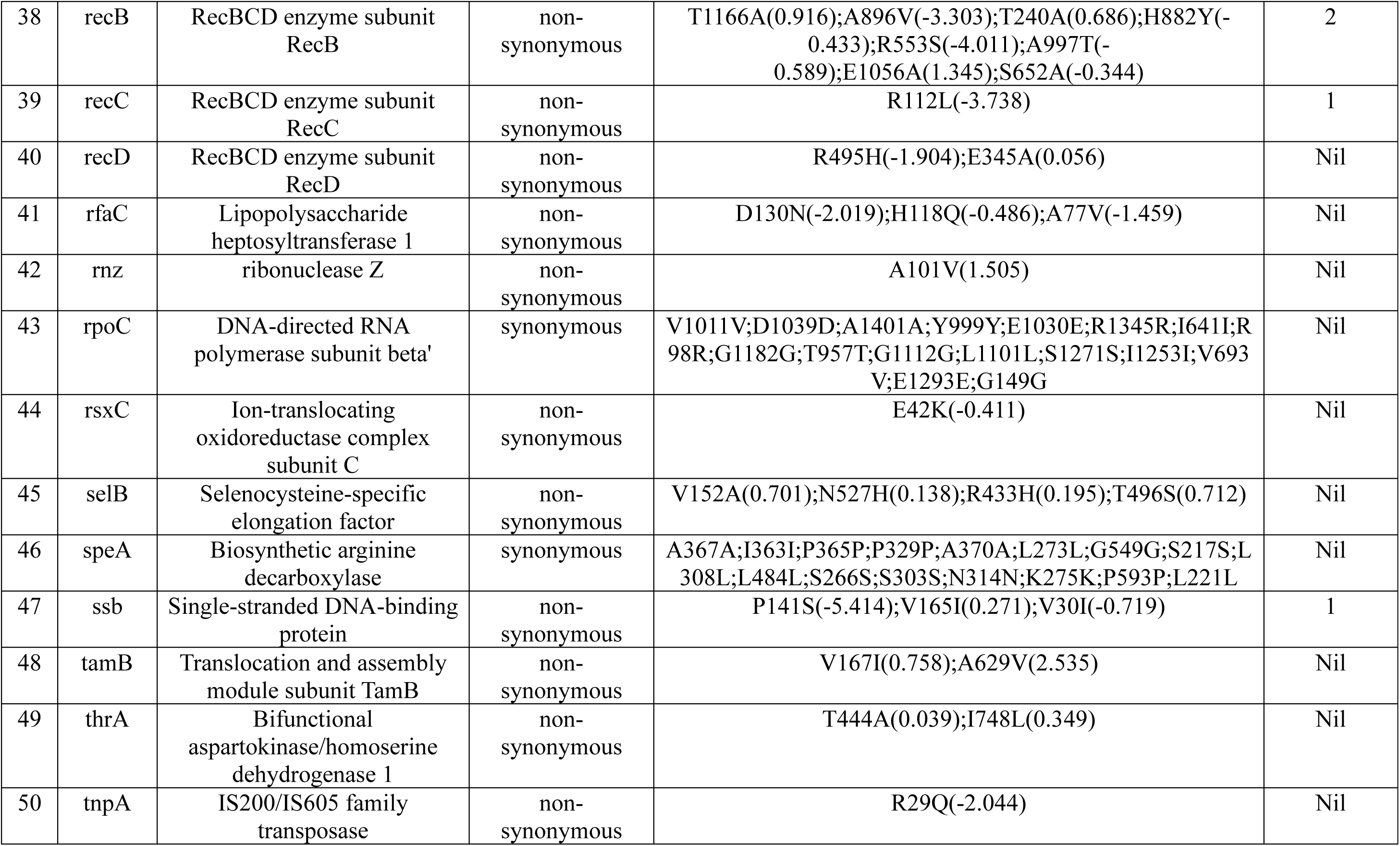
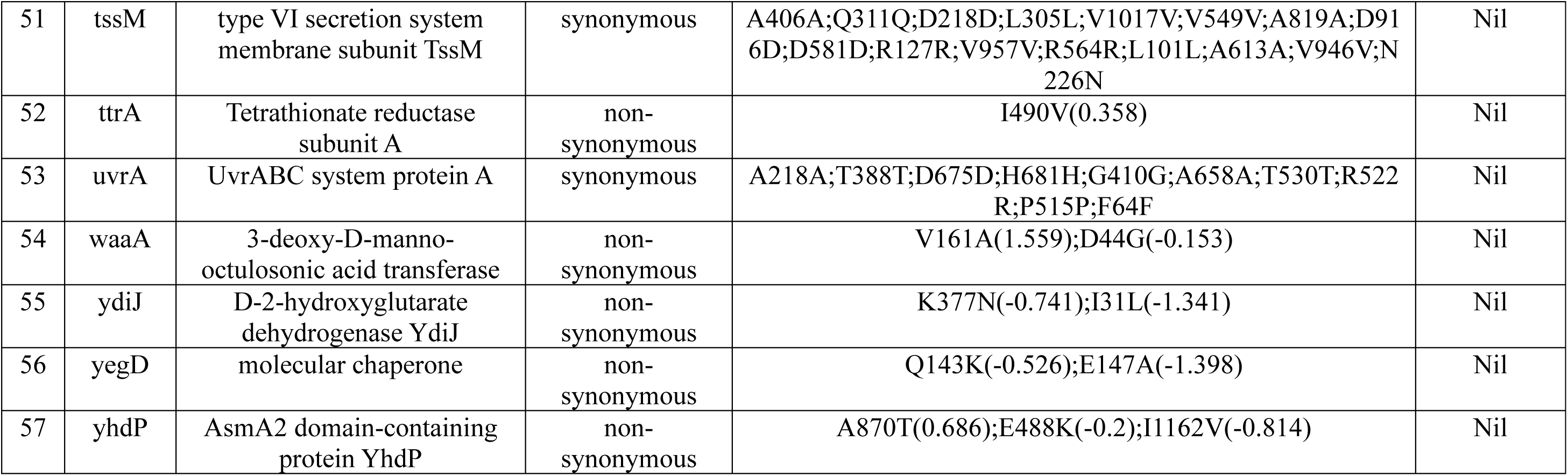
Amino acid substitutions and PROVEAN-based functional impact prediction of top variants identified in *P. mirabilis* FFPR3MR5. For each variant, the specific amino acid change and corresponding PROVEAN score are shown in parentheses. Variants with PROVEAN scores ≤ −2.5 were classified as deleterious, and the total number of deleterious variants per gene is indicated in the final column.

## Discussion

The genomic profiling of *P. mirabilis* FFPR3MR5, isolated from a fermented food source, revealed the coexistence of AMR and virulence determinants, raising potential food safety concerns. CARD analysis identified four resistance genes associated with antibiotic inactivation and efflux, conferring resistance to multiple antibiotic classes, including β-lactams, fluoroquinolones, tetracyclines, and macrolides. Similar multidrug resistance profiles have been reported in clinical *P. mirabilis* isolates, indicating that even food-derived strains can serve as reservoirs of clinically relevant AMR traits (12, 13). The presence of β-lactam and fluoroquinolone resistance determinants suggests possible selective pressure from antimicrobial exposure or horizontal gene transfer in the fermentation environment (14, 15). In addition to AMR, the strain harboured 35 virulence-related genes, including those linked to adherence (fimbriae), motility (flagella), biofilm formation, and immune evasion. These factors are essential for environmental persistence and host colonization (16, 17). The detection of capsule and stress survival genes further suggests the strain’s ability to withstand harsh conditions during food processing and storage. PHI database analysis revealed that most virulence genes were annotated as “reduced virulence” rather than “loss of pathogenicity,” implying a retained capacity for opportunistic infection. Given its close genetic relationship (99.39% ANI) with a clinical *P. mirabilis* reference genome, FFPR3MR5 may represent a lineage capable of existing across both food and clinical niches. These findings emphasize the One Health relevance of monitoring AMR and virulence in foodborne bacteria to mitigate the risk of resistance gene dissemination and safeguard food microbiological safety (18, 19).

The isolation of *P. mirabilis* strain FFPR3MR5 from a fermented soybean product underscores an emerging concern regarding the potential for opportunistic pathogens to persist and evolve within fermented food ecosystems. Traditionally, *P. mirabilis* is recognized as a commensal of the intestinal tract and an opportunistic pathogen associated with urinary tract infections, wound infections, and bacteraemia, particularly in healthcare settings (16, 17). Its detection in a fermented food matrix indicates environmental adaptability and raises questions about its transmission dynamics and virulence modulation outside clinical niches.

Comparative genomic analysis revealed that strain FFPR3MR5 shares 99.39% ANI with the clinical reference strain *P. mirabilis* (NC_010554), suggesting close evolutionary relatedness and the possible conservation of core virulence determinants (20). However, the presence of distinct single-nucleotide polymorphisms (SNPs) and deleterious variants in genes such as *bcsA, cpxA, hpmA, recB*, and *metH* points toward niche-specific genetic diversification. Many of these genes contribute to stress tolerance, biofilm formation, and DNA repair, traits advantageous for persistence in complex food fermentation environments (21). The observed mutations may represent adaptive responses to selective pressures such as low pH, high osmolarity, and microbial competition inherent to the fermentation process (22).

While the isolate’s virulence gene profile included classical determinants, such as fimbrial adhesins, flagellar components, hemolysin, and capsule-associated loci, functional predictions indicated a predominance of metabolic and environmental response genes over overt pathogenicity factors. This pattern suggests potential attenuation of acute virulence in favor of environmental survival (23, 24). Nevertheless, the co-occurrence of antibiotic resistance genes (e.g., β-lactamase, efflux transporters) and virulence-associated loci poses a latent risk. If such strains persist in food production environments or colonize consumers, they could act as reservoirs of AMR determinants transferable to commensal or pathogenic gut bacteria (25, 26).

The possibility of *P. mirabilis* being transmitted through fermented foods and contributing to community-acquired infections cannot be entirely dismissed. Although fermentation generally suppresses pathogens through acidification and microbial antagonism, occasional contamination or survival of tolerant strains could permit low-level persistence (27). Reports of *P. mirabilis* detection in dairy and fermented fish products further support its ability to adapt to non-clinical environments (4, 28). Given that *P. mirabilis* exhibits high motility, quorum sensing, and biofilm capabilities, these traits may facilitate its colonization of food-processing surfaces and subsequent reintroduction into food chains (29).

From a public health perspective, the detection of *P. mirabilis* in fermented soybean emphasizes the importance of microbiological quality control and metagenomic surveillance of traditional fermentation systems. Although FFPR3MR5 shows potential adaptation to a food-associated niche, the maintenance of virulence and resistance gene repertoires suggests that under favorable conditions, such as immune suppression or gut dysbiosis, these isolates could revert to a pathogenic state. Therefore, monitoring and genomic risk assessment of opportunistic enteric bacteria in artisanal and industrial fermented foods are crucial to prevent potential future outbreaks (30, 31).

## Conclusion

This study presents the first genomic characterization of the *P. mirabilis* strain FFPR3MR5, which was isolated from a fermented soybean product. The findings offer insights into the strain’s adaptive potential and public health significance. The genome analysis revealed a mix of virulence and AMR determinants commonly associated with clinical isolates, as well as niche-specific mutations that suggest adaptation to the fermentation environment. Comparative genomic and variant analyses showed that FFPR3MR5 is closely related to the clinical reference strain *P. mirabilis* (NC_010554), but it contains harmful substitutions in genes related to biofilm formation (*bcsA*), stress sensing (*cpxA, kdpD*), metabolism (*metH, gcvP*), virulence (*hpmA*), and DNA repair (*recB, recC, ssb*). These genetic variations may indicate adaptive responses to environmental pressures, such as pH, osmotic stress, and microbial competition that are typical in fermented food ecosystems. The coexistence of AMR and virulence genes in this food-derived isolate raises concerns that fermented foods could act as reservoirs and vehicles for opportunistic pathogens with clinical significance. Although fermentation generally inhibits pathogen growth, the persistence of *P. mirabilis* highlights the need for vigilance in both artisanal and industrial production processes.

Continuous genomic surveillance, strict hygiene practices, and risk-based monitoring of fermented foods are essential to mitigate the emergence and transmission of multidrug-resistant *P. mirabilis* and related Enterobacteriaceae. Overall, this work emphasizes the One Health approach, which integrates genomic risk assessment into food safety frameworks. It bridges environmental, food, and clinical microbiology to prevent potential future outbreaks associated with opportunistic pathogens in traditional fermented foods.

## Materials and Methods

### Sample Collection & Isolation

Fermented soybean was collected from a traditional fermented soybean producer located at Moirang, Manipur, India (24°29’38.4"N 93°46’38.5"E) on 4^th^ June 2022. The sample was transported to the laboratory under cold-chain conditions. The culture isolate was isolated on MRS agar (pH 5.5±0.2) supplemented with 0.05% cystine-HCl and 1% CaCO_3_ at 30℃ for 48-72 h in an O_2_-depleted, and CO_2_-enriched environment using an anaerobic jar with an anaerobic sachet (113829, Merck).

### DNA Extraction and identification

Genomic DNA from the washed pellets was isolated using the protocol described by Romi *et al*., 2015 (32) with additional modifications. Briefly, the washed cell pellet was resuspended with 200µl of 10mM Tris-HCl (pH 8.0) (ThermoScientific Cat. No. AFF-J22638-AE). 5 µl of lysozyme (1000 U/µl) and 2µl mutanolysin (5U/µl) were added to the cell suspension and incubated at 37°C @ 800 rpm for one h in thermoMixer®C (S/N. 5382KJ040906, Eppendorf). 20µl proteinase K(20mg/ml) was added, incubated at 55°C @ 800 rpm for 30mins in the thermoMixer®C. After proteinase K treatment, 100 mg of 0.1mm garnet beads was added and incubated at 95°C at 1500 rpm in a thermomixer. Centrifuged the tube at 10000xg for 10 min @ 4°C. Transferred 150 µl of supernatant to a fresh micro-centrifuge tube; added 1.8X volume of AMPureXP Bead (Beckman Coulter, Product No. A63881) and pipetted thoroughly. The mixture was incubated at room temperature (RT) for 10 mins. The solution was spun down and placed in the magnetic tube stand for 2-3 mins or until the solution became clear. The supernatant was discarded, and the beads were washed using freshly prepared 80% molecular-grade ethanol. Beads were air-dried for 2-3 min, or until they lost their lustre. The tube was removed from the magnetic stand. 50 µl of 10 mM Tris-HCl was added, the mixture was thoroughly pipetted, and the mixture was incubated at RT for 10 min. The tube was spun down and placed in the magnetic tube stand for 2-3 min. 48 µl of the eluted supernatant was transferred to a fresh microcentrifuge tube. The eluted DNA was quantified using a NanoDrop One (S/N AZY2123154; Thermo Fisher Scientific, USA). 16S rRNA gene was amplified using forward primer 27F-YM (5’- AGAGTTTGATYMTGGCTCAG-3’) and reverse primer 1492R (5’-TACGGYTACCTTGTTACGACTTT-3’) in 25 µl reaction volume. The reaction includes 0.2µM of each primer, 30ng DNA template, 1X Taq A buffer, 200 µM dNTPs, 1.25U Taq polymerase (KAPA Taq) (KAPA, Cat No. KK1016). The PCR cycling conditions are as follows: initial denaturation at 94℃ for 10 min; denaturation at 94℃ for 1 min; annealing at 55℃ for 1 min; extension at 72℃ for 30s for 35 cycles; final extension at 72℃ for 10 min and hold at 4℃. The amplified PCR product was sequenced on 3500 Genetic Analyzer (S/N 33135-100, Thermofisher Scientific) using the manufacturer’s protocol. The sequences were assembled in DNA Baser Assembler v5.21.0. The assembled 16S rRNA gene sequence (1443 bp) was taxonomically identified using BLASTN (33) against the NCBI rRNA type strains/16S ribosomal RNA database. Default parameters were applied, and only hits with ≥ 99% sequence identity and 100% query coverage were considered for species-level assignment. The assembled sequence was deposited in the NCBI’s GenBank.

### Whole Genome Sequencing

Sequencing library was prepared using the Nextera XT DNA library Prep kit (Cat. FC-131-1096, Illumina). 1 ng of input DNA was fragmented enzymatically and tags the DNA adapter sequences during the tagmentation step. Dual indexed adapters, Nexteara XT Index Kit V2 set A (Cat. FC-131-2001) was amplified through 12 cycle PCR. The indexed DNA library was cleaned, normalised, and denatured. 600µl of 8 pM denatured DNA library was loaded into the Illumina’s MiSeq and sequenced in 2x150bp paired-end mode to generate .fastq files.

### Genome Assembly & Annotation

Read quality was assessed using FastQC v0.12.1 (34). Low-quality reads were filtered using Trim_galore v0.6.10 (35). The genome was assembled *de novo* using SPADES v3.13.1 (36). The quality of the assembled genome was assessed with QUAST v5.0.2 (37). Genome annotation was performed in the NCBI Prokaryotic Genome Annotation Pipeline (PGAP) v6.10 (38). Genome quality was assessed with CheckM2 v1.1.0 (39), and coverage was calculated using BWA v0.7.17-r1188, SAMtools v1.3.1, and BEDTools genomecov v2.27.1 (40–42). Genome annotation for antimicrobial resistance genes, virulence genes, and pathogen-host interaction genes was performed using ABRicate, integrating against databases such as CARD (43), VFDB (44) and PHI database (45). COG and KEGG functional annotation of the genome was performed on Eggnog mapper v2.2.13. Circular genome map was generated in Proksee (46). Default parameters were used for all software unless otherwise specified.

### Phylogenetic and Comparative Genomics

Core genome alignments of *P. mirabilis* isolates were generated using Parsnp v1.2 (Harvest Suite) (47). Our assembled *P. mirabilis* FFPR3MR5 genome was used as the reference, and 101 publicly available *P. mirabilis* genomes were retrieved from NCBI RefSeq for comparative analysis.

Parsnp produced an XMFA formatted alignment (parsnp.xmfa). This file was converted into a standard FASTA multiple-sequence alignment (core_alignment.fasta) using HarvestTools v1.3 with the -x export function. The converted alignment was then used for downstream phylogenetic inference.

Maximum-likelihood phylogeny was reconstructed using IQ-TREE v3.0.1(48), where the best-fit substitution model, GTR+F+I+R7, was selected according to the Bayesian Information Criterion (BIC). Branch support was assessed using 5,000 ultrafast bootstrap (UFBoot) replicates to ensure high confidence in tree topology.

The resulting Newick tree was uploaded to the Interactive Tree of Life (iTOL) v7.2.2 (49) for visualization, circular layout rendering, annotation of bootstrap values, and customization of branch colors and labels.

### Variant call

Variants were called using FreeBayes v1.0.2 (50) from read alignments against the reference genome, enabling haplotype-based detection of single-nucleotide variants and small insertions/deletions. Variants were annotated in R using *VariantAnnotation* v1.54.1 and *GenomicRanges* v1.60.0, with gene models generated from the reference genome GFF file using *GenomicFeatures* v1.60.0 and *txdbmaker* v1.42.1, The reference genome of *P. mirabilis* (NC_010554) was indexed and loaded as a FASTA file using the *Rsamtools* package. Variant positions were extracted from the VCF file as genomic ranges, and multiallelic sites were expanded so that each alternative allele was analyzed independently. Coding consequences of variants were then predicted using the predictCoding() function with a transcript database (TxDb) constructed from the reference genome annotation. The reference genome sequence was provided as a FASTA file to enable accurate codon translation. For each variant, the corresponding amino acid change, codon alteration, and affected gene were identified and used for downstream functional impact analysis and visualization using ggplot2 v4.0.1 and the tidyverse v2.0.0 packages.

Top variant genes were selected using a variant quality threshold of 1000. The effect of amino acid changes in the genes was predicted in the Protein Variation Effect Analyzer (PROVEAN) v1.1.

## Acknowledgments

M.G.S thanks the Indian Council of Medical Research (ICMR), New Delhi, India, for providing the Junior Research Fellowship No.: 3/1/3/JRF-2019/HRD-070(31035). The Council of Scientific and Industrial Research (CSIR), New Delhi, and the CSIR-NEIST, Jorhat, Assam, are gratefully acknowledged for providing the necessary facilities, thereby facilitating the smooth progression of the study. The authors also thank the Publication & Intellectual Property Rights Committee, CSIR-NEIST, Jorhat for approving the manuscript for publication (Manuscript No: CSIR-NEIST/PUB/2025/230, dated 05-01-2026).

## Author contributions

**Romi Wahengbam,** Conceptualization, Data curation, Formal analysis, Funding acquisition, Investigation, Methodology, Project administration, Software, Resources, Supervision, Writing – original draft, Writing – review & editing | **Moirangthem Goutam Singh**, Conceptualization, Data curation, Formal analysis, Investigation, Methodology, Software, Visualization, Writing – original draft, Writing – review & editing

## Data availability

This Whole Genome Shotgun sequencing project has been deposited in DDBJ/ENA/NCBI GenBank under the accession no. JBTBYN010000000. The genome assembly version described in this paper is version JJBTBYN010000000. All raw reads were deposited in the NCBI’s Sequence Read Archive under the accession no. SRR35984731. This project is associated with BioProject PRJNA1295091. The 16S rRNA gene sequence was deposited in NCBI GenBank under the accession no. GenBank accession no. PX688172

## Supplementary tables

**Table S1:** The identity value of virulence analysis of the strain FFPR3MR5 in the VFDB database.

**Table S2:** The identity value of Pathogen host interaction analysis of the strain FFPR3MR5 in the PHI-database.

**Table S3:** The identity value of resistance gene analysis of strain FFPR3MR5 in the CARD database.

